# Germline *BRCA1* Mutation Deregulates Irradiation–Induced Cell Cycle Checkpoints in Human Mammary Luminal Progenitors

**DOI:** 10.64898/2026.08.11.743512

**Authors:** Abhijith Kuttanamkuzhi, Bowen Zhang, Kelvin Yeung, Matthew Waas, Pirashaanthy Tharmapalan, Soumili Sarkar, Curtis W. McCloskey, Olivia Drummond Guy, Hui Fang, Shreya Mishra, Thomas Kislinger, Hal K. Berman, Shane M. Harding, Rama Khokha

**Affiliations:** Princess Margaret Cancer Center, University Health Network, Toronto, ON, CA; Department of Medical Biophysics, University of Toronto, Toronto, ON, CA; Departments of Radiation Oncology and Immunology, University of Toronto, Toronto, ON, CA; Department of Laboratory Medicine and Pathobiology, University of Toronto, Toronto, ON, CA; Ahammune Biosciences, Pune, IND; Department of Computational and Systems Biology, University of Pittsburgh, Pittsburgh, PA, US

**Author notes:** Denotes equal contribution (Abhijith Kuttanamkuzhi, Bowen Zhang).

## Abstract

Mammary ductal tree comprises two epithelial lineages, basal and luminal, which harbour stem and progenitor cell subsets essential to breast physiology and are cancer-initiating precursors. Dysregulated cellular stress response is an early event in tumorigenesis, yet lineage-rooted study of primary breast epithelium pertinent to familial mutation carriers is limited. Here, we establish a lineage-resolved monolayer culture system that enables propagation of primary human mammary epithelial populations while preserving lineage identity across passages. We identify intrinsically distinct cell cycle response of luminal progenitors and basal cells to irradiation stress where the former engages both G1/S and G2/M checkpoints to achieve cell cycle arrest and the latter relies on the G2/M checkpoint. These lineage distinctions are diminished in germline *BRCA1* mutation carriers with an attenuated G1/S checkpoint in luminal progenitors. Paired transcriptomic-proteomic profiling of acute irradiation response uncovers *BRCA1+/−* luminal progenitor population with sustained AKT-mTOR signaling and compromised cell cycle arrest marking an early deviation in the stress response of these purported cells-of-origin of aggressive breast cancers known to arise in *BRCA1* germline mutation carriers. Our study provides a functional framework for determining critical events in the expansion of genomically altered mammary epithelial cells in the high-risk breast to enable future preventive interventions.

## Introduction

Germline *BRCA1* mutation carriers are particularly vulnerable to developing breast cancer with up to 85% lifetime risk^1–3^. Routine screening including mammograms and magnetic resonance imaging in conjunction with prophylactic mastectomy is the only effective preventative intervention for this high-risk demographic^4^. Understanding the cellular alterations induced by *BRCA1* heterozygosity will provide insights into early breast tumorigenesis that could be leveraged for less invasive risk reduction strategies.

Cell-of-origin studies have proposed that each PAM50-defined molecular breast cancer subtype corresponds to distinct basal and luminal cell populations within the mammary epithelium^5–8^. *BRCA1* mutations carriers are predisposed towards developing the aggressive basal-like subtype, typically lacking receptors for ovarian hormones and HER2, at a younger age prior to menopause. Genetically engineered mouse models and single-cell transcriptomics studies have provided evidence that basal-like breast cancers arise from the luminal progenitor (LP) population, which is expanded in *BRCA1* mutation carriers (*BRCA1+/−)*^9,10^. The signaling mechanisms that regulate mammary stem/progenitor cell proliferation and survival are often implicated in breast cancer development^11^. However, there are currently limited ways to study how aberrant molecular phenotypes are reflected in the behaviour of primary mammary epithelial cells of germline *BRCA1* mutation carriers.

Direct interrogation of FACS purified epithelial populations has shown distinctions in lineage intrinsic capacity to handle cellular stress. The LP population has enhanced mitochondrial metabolism, more robust DNA damage repair capacity and antioxidant defense than basal cells^12–14^. These observations suggest that LP cells may be more adept at surviving following genotoxic insults. How the *BRCA1* heterozygous state impacts lineage-specific molecular stress response remains currently unknown due to a paucity of suitable *ex vivo* models. Existing methods for culturing primary human mammary epithelial cells as monolayers are not lineage-resolved and reliant on stromal feeder cells^15–18^. We developed a workflow to propagate breast-derived basal and LP cell populations over multiple generations while maintaining their lineage identity. Using this monolayer culture system, we identified lineage-specific cell cycle response along with its molecular underpinnings, upon gamma irradiation. Moreover, our study revealed the altered stress response of *BRCA1+/−* LP cells, shedding light on early maladaptive events in the epithelial cell population purported to be the origin of the most aggressive breast cancer subtype.

## Results

### Breast-derived single-lineage monolayers retain lineage features

We devised a new 2D feeder-free cell culture method guided by morphology, flow cytometry profiling, and multi-OMICs to establish and validate lineage-specific human mammary epithelial monolayers. To control for reproductive cycle effects, we selected eight human breast tissue samples histologically confirmed to be premenopausal and in the follicular phase of the menstrual cycle (Age range: 37 to 52 yrs). Four samples possessed pathogenic germline mutations in a single *BRCA1* allele (*BRCA1+/−*), while the other four lacked any known high-risk germline mutations (non-carriers) (Figure S1A). We purified basal (CD49f^hi^/EpCAM^lo-mid^), luminal progenitor (LP: CD49f^hi^/EpCAM^hi^), and mature luminal (ML: CD49f^lo^/EpCAM^hi^) cell populations from the viably cryopreserved epithelium-enriched fractions^19,20^ of these donors by fluorescence activated cell sorting (FACS) to generate monolayers (Figure 1A). The basal and LP populations attained distinct morphology resembling that of basal and luminal colonies from standard mammary colony formation assays, respectively (Figure 1B). These single-lineage monolayers were optimally passaged at 70% confluency every 2 to 4 days for up to 7 generations, after which proliferative capacity was greatly reduced. The ML population, enriched in more differentiated luminal cells, had limited proliferative capacity as monolayers and could not be passaged beyond 3 generations even with TGFß inhibitors^21,22^. Harvested LP and basal monolayer cells maintained the lineage-specific EpCAM/CD49f staining profiles in flow cytometry analysis (Figure 1C). All downstream analyses were performed with basal and LP monolayers derived from the same eight donor samples. Recognizing the utility of 3D culture models, cell suspensions generated from the epithelium-enriched fraction of breast tissues were also cultured as organoids with basement membrane extract using the reported 3D on-top method and defined serum free media^23–25^. Non-carrier and *BRCA1+/−* organoid cultures maintained all three epithelial populations (basal, LP, ML) for at least 3 generations (Figure 1D) and produced polarized spheroidal structures with basal cells on the exterior as previously described (Figure 1E).

**Figure 1.**
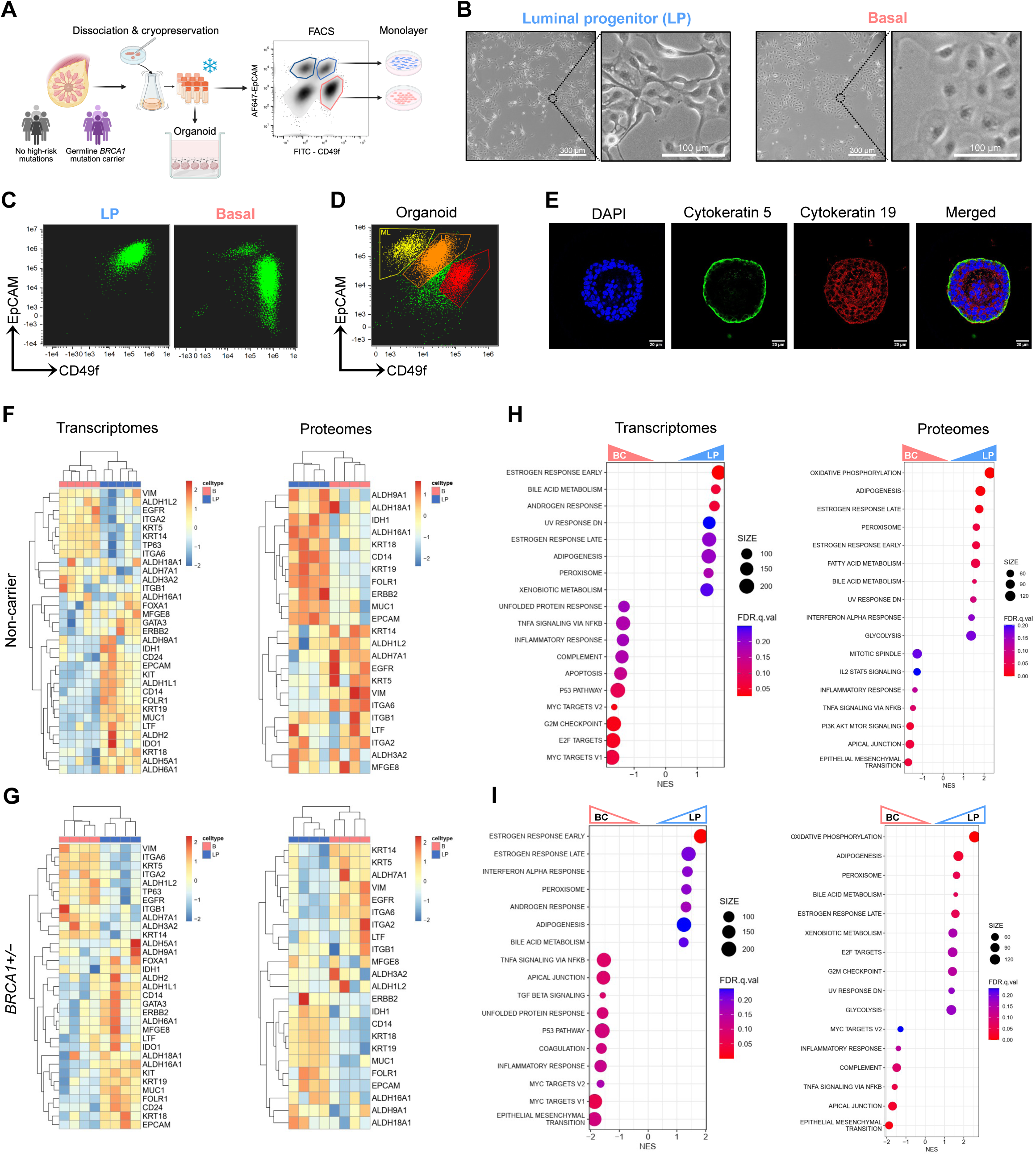
Breast-derived single-lineage monolayer cultures retain lineage features. A) Schematic of breast tissue collection and processing workflow leading to FACS purification of specific mammary epithelial populations. B) Representative images of LP and basal monolayer cultures at passage 3 (Scale-300 µm) and C) corresponding flow cytometry profiles. D) Flow cytometry profiles of organoid cultures showing lineage marker expression. E) Representative cross-sectional of organoid culture after 16 days of culture, immunostained for lineage markers. Heatmaps showing expression of lineage-specific markers at both RNA and protein levels in lineage-resolved monolayer cultures derived from F) non-carrier and G) BRCA1 mutation carrier breast tissues. Transcriptomic and proteomic-driven GSEA using Hallmark gene sets showing top 10 differentially enriched pathways between LP and basal monolayers from H) non-carriers and I) BRCA1 mutation carriers.

Non-carrier monolayers displayed cytokeratin expression expected of their lineage (Basal: KRT5, KRT14; LP: KRT18, KRT19) along with other known markers based on published lineage-defining gene sets at both transcript and protein levels^12,26^ (Figure 1F, S1B). Gene set enrichment analysis (GSEA) performed on bulk transcriptomics identified androgen and estrogen response as top enriched terms in non-carrier LP monolayers, and the latter was also found enriched at the protein level (Figure 1H). p53 pathway and inflammatory response-associated terms were enriched in basal monolayers at both transcript and protein levels (Figure 1H). Proteomics analysis additionally revealed enrichment of Oxidative Phosphorylation and Fatty Acid Metabolism terms, in non-carrier LP monolayers recapitulating lineage features previously identified (Figure 1H). *BRCA1+/−* monolayers also retained distinct morphology and lineage-marker expression (Figure 1G, S1C, D). Parallel GSEA analyses of *BRCA1+/−* transcriptomics and proteomics revealed comparable pathway enrichment trends to non-carriers (Figure 1I), indicating normal characteristics under baseline culture conditions. Together, monolayers and organoids, serve as complementary *ex vivo* models for investigating the effect of germline *BRCA1* mutations on DNA damage response in a lineage-specific manner.

### Irradiation-induced lineage distinctions in cell cycle dynamics are attenuated in *BRCA1+/−*

To assess lineage-specific DNA damage responses, LP and basal monolayers were exposed to 10 Gy gamma irradiation and tracked over 120 hours by live cell imaging using a fluorescent nuclear probe (SPY505-DNA) at 2-hour intervals (Figure 2A). Proliferation curves, represented as fold change to initial cell count, showed that unirradiated *BRCA1+/−* monolayers had higher proliferation rates than non-carrier counterparts (Figure S2A) with basal monolayers displaying greater proliferation than LP monolayers for both genotypes (Figure S2A, B). Irradiation drastically suppressed proliferation of both basal and LP monolayers without inducing substantial cell death for approximately 100 hours. Non-carrier basal cell count fold change fell below 1 after 100 hours, indicating net cell death. Interestingly, after an initial decrease within the first 24 hours, non-carrier LP cell numbers remained stable over the next 100 hours, indicating growth arrest (Figure 2B). Image analysis showed significantly more metaphase cells in non-carrier basal monolayers than LP monolayers at 90 hours post-irradiation (Figure 2D, Supplementary Video 1-8), suggesting that irradiated basal cells proceeded through mitosis and later underwent cell death. Unlike non-carriers, *BRCA1+/−* basal and LP monolayer proliferation curves remained similar between 25 and 75 hours post irradiation (Figure 2C). Additionally, LP cells carrying a *BRCA1* mutation displayed a higher number of mitotic events 90 hours post-irradiation, diminishing a prominent non-carrier lineage distinction (Figure 2D).

**Figure 2.**
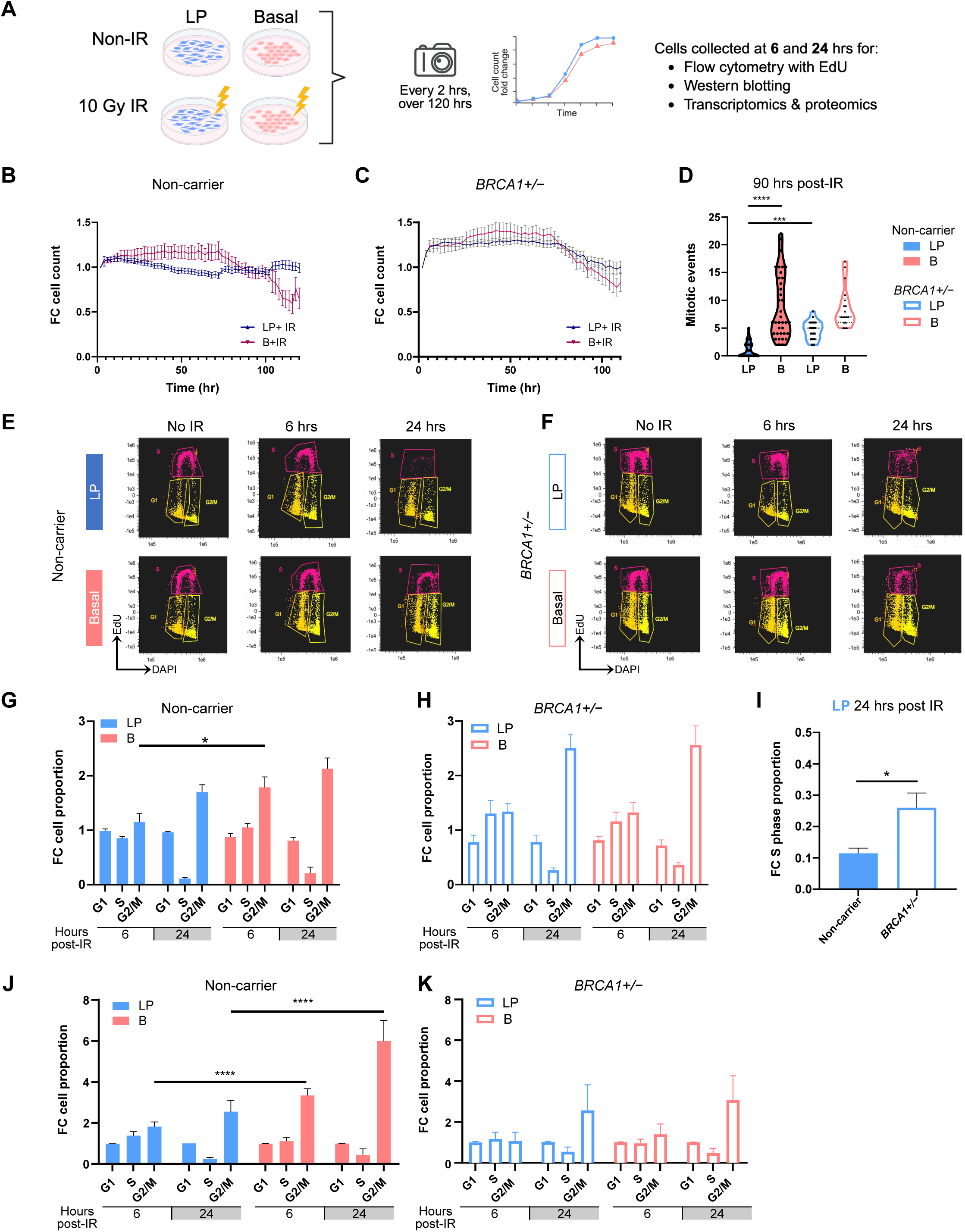
Irradiation-induced lineage distinctions in cell cycle dynamics are attenuated in *BRCA1+/−*. A) Experimental workflow used for monolayer irradiation-stress response assays. Growth curves show fold change (FC) in nuclei count over 120 hours with images captured at every 2 hours showing proliferation rate of irradiated (10 Gy) B) non carrier and C) BRCA1+/− LP and basal cells. D) violin plot showing the number of mitotic events per frame in LP and basal cells post-irradiation (N=4, Mann-Whitney test). Representative flow cytometry plots showing cell cycle phase distribution of E) non-carrier and F) BRCA1+/− cells measured by EdU incorporation and nuclear content (DAPI) in monolayer cells at 6 and 24 hours post-irradiation. The fold change shift of cell cycle phase proportions relative to non-irradiated control (non-IR) in G) non-carrier and H) BRCA1+/− monolayers at 6 and 24 hours post-irradiation. I) Bar plot showing fold change in proportion of S-phase cells 24 hours post-irradiation in LP cells of non-carriers compared to BRCA1 mutation carriers (derived from G and H). Fold change in cell cycle phase proportions of irradiated organoid cultures relative to non-IR of J) non carrier and K) BRCA1 mutation carriers determined by EdU incorporation and DAPI staining. (N=4, Two-way ANOVA with Sidak’s multiple comparisons Statistical analysis was performed using tests as mentioned. *P < 0.05, **P < 0.01, ***P < 0.001and ****P < 0.0001).

We then performed EdU labeling in monolayers at 6 and 24 hours post-irradiation to capture early cell cycle kinetics that could underlie later proliferation patterns. Non-carrier LPs showed a slight decrease in S phase cells at the 6-hour time point, which was drastically reduced by 24 hours (Figure 2E, G). On the other hand, non-carrier basal cell cycle profiles indicated reduced but continued cycling 6- and 24-hours post irradiation (Figure 2E) consistent with their proliferation curves. To account for interpatient variation of breast tissue samples, cell cycle phase proportions were normalized to corresponding unirradiated controls. At the 6-hour time point, non-carrier basal monolayers displayed a significantly greater increase in G2/M cells than matched LP monolayers (Figure 2G). By 24-hours post-irradiation, non-carrier LP monolayers had a sharper decrease in S-phase cells accompanied by an increase in the G2/M fraction consistent with cell-cycle arrest (Figure 2G). In contrast, the cell cycle phase shifts of *BRCA1+/−* LP monolayers were more akin to their basal counterparts at both 6 and 24-hours (Figure 2F, H). Specifically, *BRCA1+/−* LPs maintained a higher proportion of S-phase cells at 24 hours, possibly due to a loosened G1/S checkpoint (Figure 2I). Lineage-specific cell cycle perturbations induced by 10 Gy irradiation were recapitulated in non-carrier organoids and diminished in *BRCA1+/−* organoids (Figure 2J, K). These data suggest that lineage-intrinsic cell cycle regulation of human mammary epithelial cells upon DNA damage is eroded in *BRCA1* mutation carriers.

### Irradiated *BRCA1*+/− LPs have delayed cell cycle checkpoint activation

We next assessed activation of key cell cycle regulators by Western blotting in monolayer lysates at 6 and 24 hours after irradiation. Phosphorylated H2AX was detected in both non-carrier and *BRCA1+/−* monolayers, confirming the presence of unresolved double stranded DNA breaks (Figure S3 A, B). In non-carrier LPs, phosphorylated Retinoblastoma protein (p-Rb) was already significantly reduced at the 6-hour timepoint indicating early and robust G1/S checkpoint activation. This response progressed to stress-induced cell cycle exit by 24-hour, as evident by increased p21 expression and decreased Cyclin D1 levels^27,28^ (Figure 3A, B). Reduction of p-Rb in irradiated *BRCA1+/−* LPs was, in contrast, delayed until 24-hours (Figure 3C, D). Although p21 was induced, Cyclin D1 levels were not significantly reduced in *BRCA1+/−* LPs at 24 hours post-irradiation (Figure 3D). A significant increase in phosphorylated p53 (Ser15) at 6 hours post-irradiation was observed only in non-carrier LPs, whereas p53 phosphorylation remained relatively unchanged in *BRCA1+/−* LPs (Figure S3B). This is supportive of a relatively stronger G1/S checkpoint in LP cells relative to basal cells and attenuation of this lineage distinction in *BRCA1* mutation carriers. In comparison, p-Rb, p21 and Cyclin D1 levels were not significantly altered in irradiated basal monolayers of both genotypes relative to unirradiated controls at 24 hours post-irradiation (Figure 3B, D).

**Figure 3.**
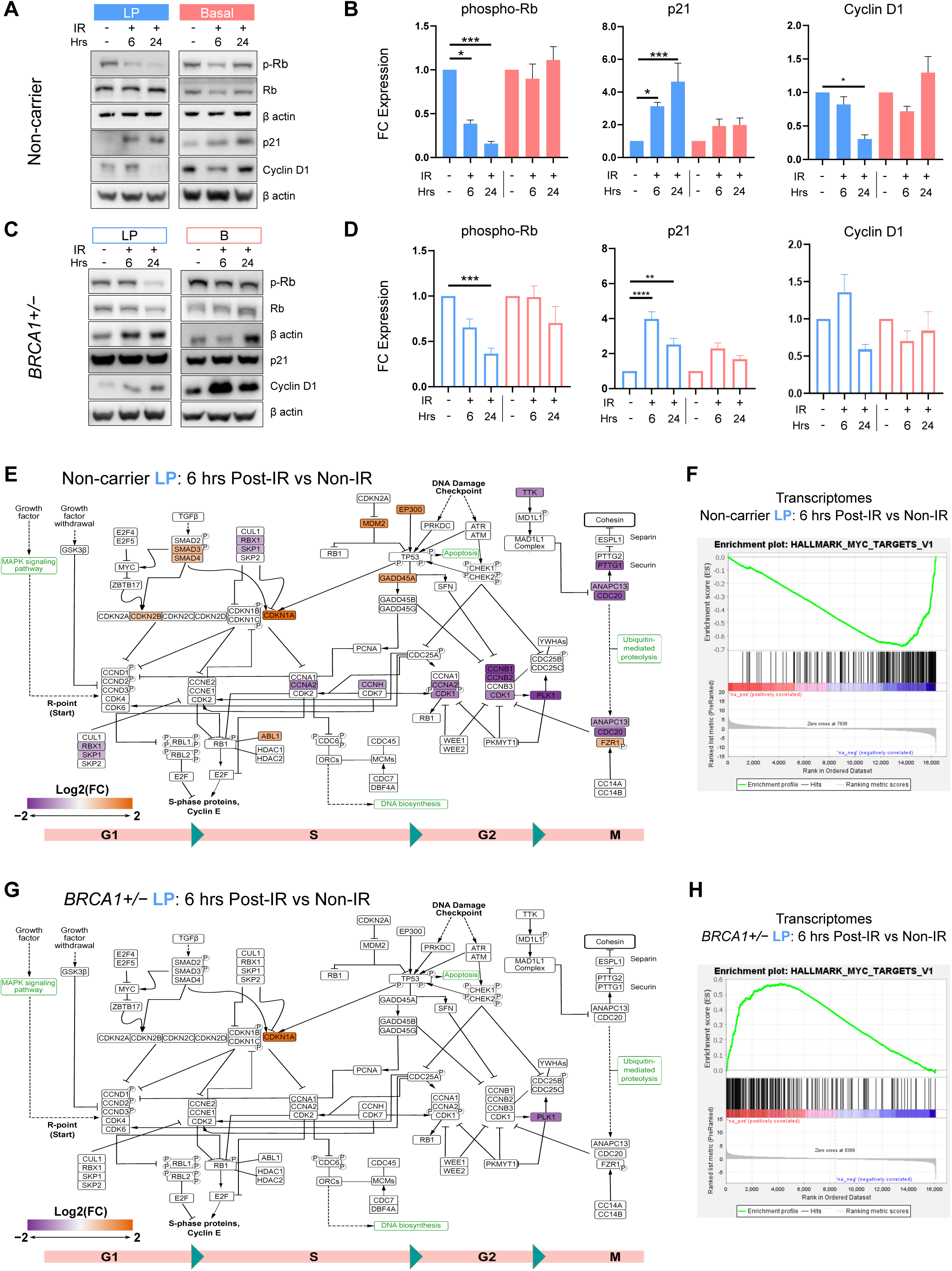
Irradiated *BRCA1*+/− LPs have delayed cell cycle checkpoint activation. Representative western blots of G1 checkpoint markers (Rb phosphorylation and p21) and Cyclin D1 in A) non-carrier and C) BRCA1+/− LP and basal monolayers with or without irradiation. Quantification of changes in these markers are expressed as fold change (FC) to matched non-IR controls for B) non-carriers and D) BRCA1+/− LP and basal monolayers (N = 4, 1-way ANOVA and Holm-Sidak’s multiple comparison). KEGG cell cycle pathway diagram showing significantly altered transcripts with a FC of at least 1.5 relative to non-IR control for E) non-carrier and G) BRCA1+/− LP monolayers 6 hours post irradiation. GSEA plot for the Hallmark MYC Targets term from transcriptomics comparing 6 hr post irradiation to non-IR control of F) non-carrier and H) BRCA1+/− LP cells.

To assess gene expression changes associated with cell cycle checkpoint signalling, we analyzed monolayer transcriptomics collected at the established time points following irradiation. Differentially expressed genes (DEGs) of non-carrier LPs at the 6-hour time point were mapped onto the KEGG Cell Cycle pathway. We observed marked downregulation of key cell cycle regulators, including Cyclin-dependent kinases (CDKs), Aurora kinase B (AuroraB) and Polo-like kinase 1 (PLK1) relative to unirradiated controls (Figure 3E). Parallel analysis performed on *BRCA1+/−* LP monolayers identified far fewer DEGs suggesting impaired cell cycle checkpoint activation associated with *BRCA1* heterozygosity (Figure 3G). Furthermore, transcriptomics-driven pathway enrichment analysis identified MYC targets to be downregulated in non-carrier LPs post irradiation, while upregulated in *BRCA1*+/− LPs (Figure 3F, H, S4A). Perturbation of MYC targets, which includes a large fraction of cell cycle and metabolic signalling regulators can rewire cell cycle checkpoints^29^. Collectively, these data demonstrate that the normally rapid post-irradiation cell cycle checkpoint engagement in non-carrier LP cells is delayed in germline *BRCA1* mutation carriers.

### Irradiated *BRCA1+/−* LP cells sustain mTOR and reduce senescence-associated signaling

To investigate the broader effects of *BRCA1* heterozygosity on the irradiation stress response, we performed lineage-resolved GSEA on matched monolayer transcriptomics-proteomics data. mTORC1 signaling emerged as one of the most highly enriched pathways in *BRCA1*+/− LPs relative to non-carrier LPs at the transcript level 6 hours post-irradiation (Figure 4A) and at the protein level 24 hours post irradiation (Figure 4B). Enrichment of the mTORC1 signaling pathway in *BRCA1+/−* basal cells was seen only at the transcript level 6-hours post irradiation and not at the protein level (Figure S4B). Additional metabolic terms including Oxidative Phosphorylation and Fatty Acid Metabolism were prominently enriched in irradiated *BRCA1+/−* LPs at both time points relative to non-carrier LPs (Figure S4A). These findings suggest deregulation of mTOR signaling and associated metabolic programs in the *BRCA1+/−* LP response to DNA damage. The downregulation of metabolic pathways in non-carrier LPs relative to *BRCA1+/−* counterparts, could also be reflective of a concomitant decrease in metabolic activity upon cell cycle arrest.

**Figure 4.**
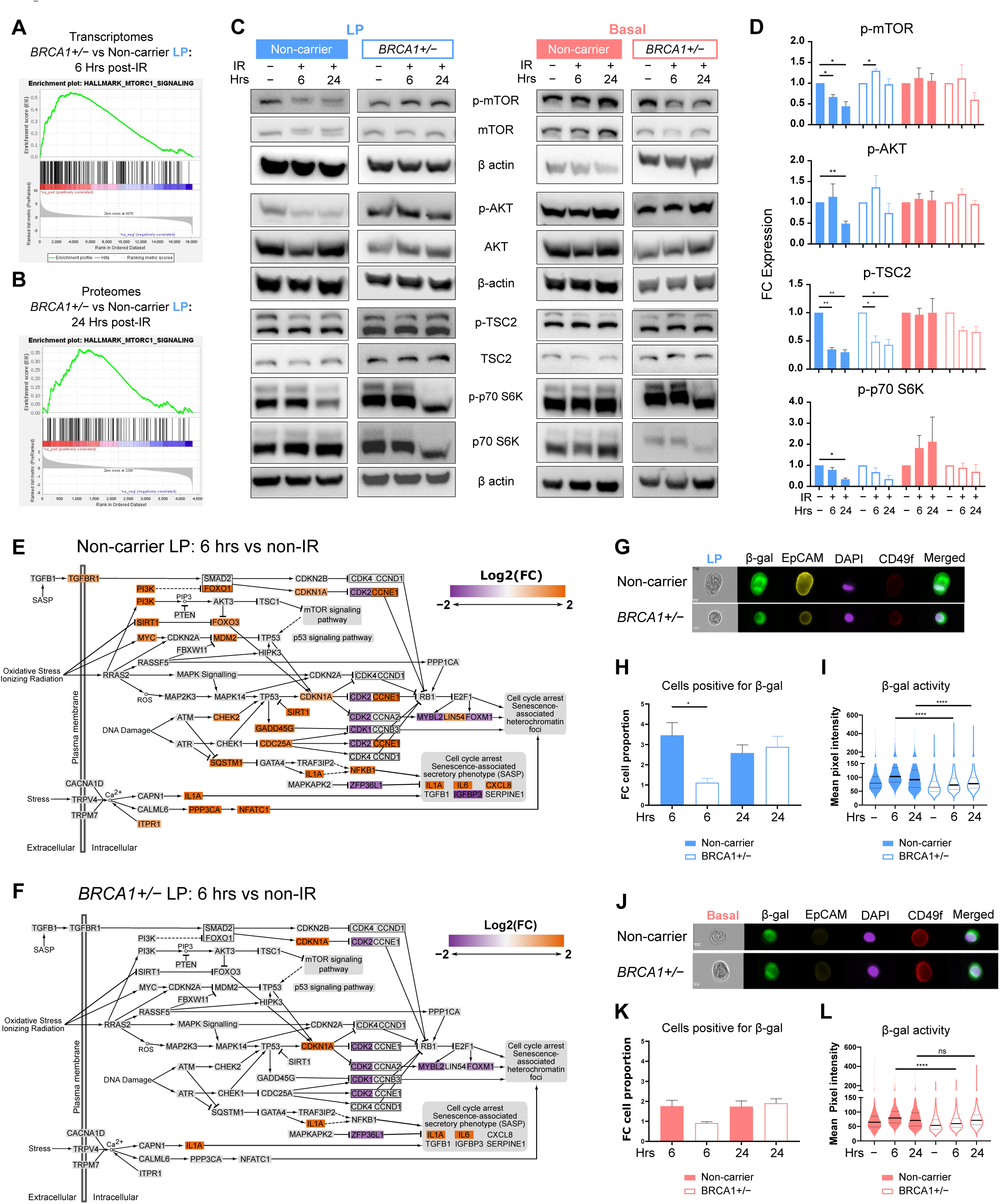
Irradiated *BRCA1+/−* LP cells sustain mTOR and reduce senescence-associated signaling. GSEA plot for the Hallmark mTORC1 signaling term from A) transcriptomics and B) proteomics comparing BRCA1+/− LP to non-carrier LP. C) Representative western blot images probed for different proteins along the AKT-mTOR pathway, along with corresponding phosphorylated protein levels and β-actin as loading control. (D) Quantification of changes in mTOR signalling pathway proteins expressed as fold change (FC) to matched non-IR controls in non-carrier and BRCA1+/− monolayers. (N=4, 1-way ANOVA and Holm-Sidak’s multiple comparison). KEGG cellular senescence pathway diagram showing significantly altered transcripts with a FC of at least 1.5 relative to non-IR control for E) non-carrier and F) BRCA1+/− LP monolayers 6 hours post irradiation. Representative images from imaging flow cytometry showing SA-β-gal activity, lineage markers (EpCAM and CD49f) and nuclei (DAPI) (Scale-10mm) in G) LP monolayers. H) Fold change in SA-β-gal positive LP monolayer cells post irradiation normalised to their respective unirradiated control. I) Violin plot showing the mean pixel intensity of SA-β-gal in LP monolayer cells after irradiation (N=3, n≥2300, Kruskal-Wallis test followed by Dunn’s multiple comparison test). Representative images from imaging flow cytometry showing SA-β-gal activity, lineage markers (EpCAM and CD49f) and nuclei (DAPI) (Scale-10mm) in J) basal cells. K) Fold change in SA-β-gal positive Basal monolayer cells post irradiation normalised to their respective unirradiated control. L) Violin plot showing the mean pixel intensity of SA-β-gal in basal monolayers after irradiation (N=3, n≥2300, Kruskal-Wallis test followed by Dunn’s multiple comparison test). Statistical analysis was performed using tests as mentioned (P < 0.05, **P < 0.01, ***P < 0.001and ****P < 0.0001).

The *BRCA1+/−* associated stress response deregulation is intriguingly more prominent in LP than basal. To investigate lineage-specific mTORC1 activity following irradiation, we performed Western blots to interrogate the mTOR signaling axis, including TSC2, AKT and S6K on monolayer cell lysates (Figure 4C, D). In non-carrier LPs, irradiation induced a significant downregulation of phosphorylated mTOR (p-mTOR) cells at both 6- and 24-hour time points. Phosphorylation of TSC2 at Thr1462 (p-TSC2), as well as activating phosphorylation of AKT at Ser473 (p-AKT), were significantly reduced, together signifying negative regulation upstream of mTOR. Activating phosphorylation of S6K (p-p70-S6K), was only significantly decreased at 24 hours, suggesting participation of additional signalling events that regulate this downstream effector. A decrease in p-TSC2 was also observed in *BRCA1+/−* LPs following irradiation. However, p-mTOR levels did not decrease, nor did p-AKT or p-p70-S6K. mTOR activity was minimally affected in basal cells of both genotypes 24 hours post irradiation, suggesting deregulation of mTOR signaling is specific to *BRCA1+/−* LPs. These findings align with emerging evidence that pre-neoplastic, BRCA1-deficient mammary luminal progenitors as well as fully transformed BRCA1-mutant breast tumor cells exhibit hyperactivation of the PI3K/AKT/mTOR pathway, underscoring the critical role of this cascade in early BRCA1-driven breast carcinogenesis^30–32^.

Transcriptomic analyses further revealed upregulation of TGFβ and TNFα signalling in non-carrier LP cells relative to basal cells at 6 hours following irradiation, which was notably absent in *BRCA1+/−* monolayers (Figure S4C, D). Enrichment of inflammatory and stress-associated pathways, together with a cell cycle arrest phenotype raised the possibility that non-carrier LPs may be initiating senescence pathways shortly after irradiation, especially in the context of primary cell cultures^33–37^. Non-carrier LPs displayed a greater upregulation of genes from senescence-associated transcriptional programs displayed on the KEGG Cell Senescence pathway map (Figure 4E) compared to *BRCA1+/−* LPs (Figure 4F) 6 hours post irradiation. We then assessed β-galactosidase (β-gal) expression and activity in irradiated monolayer cells by imaging flow cytometry. A significantly larger proportion of non-carrier LPs was positive for β-gal (β-gal+) than *BRCA1+/−* LPs 6-hours post-irradiation, relative to unirradiated control (Figure 4G, H). Moreover, β-gal+ LPs cells of non-carriers displayed significantly higher mean pixel intensity, indicating greater enzymatic activity than *BRCA1+/−* counterparts (Figure 4I). Parallel analyses in basal cells showed non-significant differences in the proportional increase of β-gal+ cells between non-carrier and *BRCA1+/−* (Figure 4J, K) at both time points, but β-gal activity was significantly increased in non-carrier basal cells (Figure 4L).

Altogether, irradiated *BRCA1*+/− LPs exhibit deregulated mTOR signaling and associated metabolic processes along with reduced progression towards a senescence-like state. These deviations from the non-carrier LP response reinforce the theme that *BRCA1*+/− LPs adopt a stress response more closely resembling that of basal cells following DNA damage. Our data demonstrates that *BRCA1* heterozygosity specifically compromises LP cellular stress response.

## Discussion

Establishing mammary lineage-specific stress response provides critical insight into why particular cell populations are prone to oncogenic transformation and how germline *BRCA1* mutations elevate breast cancer risk at the molecular level. In this study, we established a feeder-free methodology to propagate FACS-purified mammary basal and LP cell populations from human breast tissues as adherent monolayer cultures. Using integrated transcriptomic, proteomic, and flow cytometric analyses, we show that these monolayers retain canonical lineage characteristics over successive passages and shed light on the underlying molecular pathways. Surprisingly, basal and LP cells without a *BRCA1* mutation mount distinct cell cycle response when exposed to gamma irradiation. LPs robustly engage the definitive G1/S and G2/M checkpoints known to be essential for ensuring conformity of cell division in response to stress, whilst basal cells predominantly rely on G2/M checkpoint activation for cell cycle arrest. Parallel comparisons of these two primary breast lineages from germline *BRCA1*+/− mutation carriers reveal further distinctions. *BRCA1*+/− LPs have an attenuated G1/S checkpoint response and cell cycle arrest, shifting their behavior towards a basal-like response pattern. To our knowledge, this is the first study to identify how *BRCA1* heterozygosity could confer lineage-specific proliferative advantage following genotoxic stress beyond its established function in DNA damage repair^38^.

Single cell transcriptomic and proteomic studies have identified LP subsets that express basal lineage markers (Basal-luminal cells) that increase with age and are transcriptionally similar to the basal-like breast cancer subtype^39–41^. The resemblance of the *BRCA1*+/− LP irradiation stress response to that of basal cells could be indicative of lineage infidelity, which has been proposed as an early event in oncogenic transformation. Aberrant cell cycle regulation may allow for accelerated mutation accumulation or propagating existing mutations specifically in the LP population. LP cells irrespective of germline *BRCA1* mutation status demonstrate a greater tendency to persist following DNA damage compared to basal cells. This mirrors our prior observation where LP colony-forming capacity in both mouse and human systems were less impacted by poly (ADP-ribose) polymerase (PARP) inhibitors, which indirectly induce double-strand DNA breaks^13^. We propose that robust activation of cell cycle checkpoints and senescence-associated transcriptional programs^42^ seen in LP cells after irradiation are required as protective mechanisms to limit the inheritance of mutations into progeny cells.

Molecularly, our matched transcriptome-proteome pathway analyses along with biochemical validation, revealed that *BRCA1*+/− LPs sustain mTOR activity following irradiation, unlike non-carrier counterparts. On the other hand, basal cells do not modulate the AKT-mTOR axis as part of their early irradiation response. This persistent signaling likely functions as a critical survival adaptation, enabling BRCA1-haploinsufficient progenitors to bypass DNA damage-induced cell cycle checkpoints and undergo clonal expansion despite harboring high levels of genomic instability^31,32^. Given that the mTOR pathway is a key signalling hub that integrates intracellular and extracellular cues, including cellular metabolism^43–46^, its unique involvement in *BRCA1*+/− LPs may in part be related to the concomitant enrichment of metabolic processes seen in *BRCA1*+/− compared to non-carrier LPs^30^. Mammary LP and basal populations are known to have inherently distinct metabolic identities with evidence of enhanced mitochondrial metabolism as a feature of LPs^12,14^. The effect of *BRCA1* mutations on mTOR-mitochondria crosstalk warrants further investigation.

Clinically, prophylactic mastectomy combined with intensive surveillance remains the most effective risk-reduction strategy for germline *BRCA1* mutation carriers. Novel *ex vivo* cultures derived herein from primary breast specimens from high-risk women provide a tractable path to define molecular consequences of *BRCA1* heterozygosity and to identify prevention-oriented vulnerabilities. Further work dissecting the lineage-intrinsic dependence on mTOR signaling and oxidative phosphorylation may inform combinatorial interventions to target cancer-prone epithelial populations^12,26,47,48^. The lineage-preserving nature of our monolayer system offers a practical bridge between OMICs data driven hypotheses gleaned from high-risk tissue and their functional validation at the cellular levels^49–51^. Finally, the straightforward, feeder-free workflow is scalable and could enable high-throughput perturbation studies, modeling of early transformation events, or serve as a modular component for building more complex 3D culture systems. In summary, we show that *BRCA1* mutant luminal progenitors exhibit an altered DNA damage stress response and cell behaviour providing new insights into cancer-initiating precursors in germline *BRCA1* mutation carriers.

### Limitations of the study

Our analyses were focused on the effect of DNA damage on the proliferation of monolayers derived from breast epithelial populations with known clonogenic capacity. The use of 10 Gy gamma irradiation, although not physiological, has been used in experimental settings to gain insights into how cells respond to DNA damage insult. We noted considerable variation in the irradiation-induced molecular responses of *BRCA1* mutation carrier-derived monolayers. This may be attributable to inter-donor heterogeneity or different types of pathogenic *BRCA1* mutations (e.g. Frameshift, nonsense, missense). The monolayer model could be leveraged to study cell subsets with high clonogenic potential and/or augmented with ovarian hormone stimulation.

## Methods

### Human breast tissue dissociation and processing

All human breast tissues used in this study were acquired with informed consent and approval by the Institutional Research Ethics Board of the University Health Network (UHN, Toronto). Human breast tissue samples were processed following a published overnight dissociation protocol^52^. The epithelium-enriched fraction (pellet A) is thawed and digested with Trypsin and dispase to produce single cell suspensions for FACS. Epithelial fragments from pellet A are washed in ice-cold DMEM/F12 (Gibco, 11330-032) and pelleted by centrifugation. The resulting cell pellet is digested with pre-warmed Trypsin-EDTA (0.25%, STEMCELL Technologies, 07901) for 3 min and quenched with ice-chilled Hank’s Balanced Salt Solution supplemented with 2% fetal bovine serum (HF). The process is repeated following centrifugation with pre-warmed dispase (1 U/mL, STEMCELL Technologies, 07913) and 2000 U/mL DNase (Sigma, D4513). Following quenching with HF, the resulting cell suspension is passed through a 40-um filter, centrifuged and resuspended in 1 mL HF for cell counting by hemacytometer. All centrifugation steps are at 350 x g for 5 min, 4°C.

### Sorting human mammary epithelial cell populations by FACS

Primary tissue-derived epithelial cell fractions are retrieved from liquid nitrogen storage; they are thawed at 37 °C water bath. The entire contents of the vial are dumped into 10ml ice-cold DMEM in a 15ml tube. The cells are spun down at 350xg for 5 minutes at 4° C. The supernatant is removed, and the tissue fraction is dissociated with 2ml warm 0.25% Trypsin-EDTA by thorough pipetting for exactly 3 minutes. The trypsin is deactivated by adding 10 mL cold HBSS-2% FBS mix (HF). The fragments are spun down at 350g for 5 minutes. After removing the supernatant, the fragments are dissociated into single cells using 2ml dispase with 100 μL DNase (2000U/ml). Mix thoroughly for 3 minutes and inactivate the dispase by adding 10ml HF. The samples are passed through 40-micron filters to collect single cells and spun down at 350g for 5 minutes. Pellet is resuspended in 1ml PBS and are counted. Cells are then stained using 1:50 EpCAM-AF647, 1:100 CD49f-FITC, CD45-PE/Cy7 (1:200), and CD31-PE/Cy7 (1:50) for 30 minutes at 4° C. Cells are washed with 1ml PBS and spun down at 350g for 5 minutes. Dead cells were stained using DAPI (1:10,000) for 5 minutes at RT. Cells are washed and spun down again and are taken for FACS. Sorting was carried out at Princess Margaret Flow Cytometry Facility using the BD FACS ARIA Fusion. Cells were collected in 5ml flow tubes coated with HF. Cell pellets were centrifuged at 350xg for 5 minutes, and the supernatant was removed. Cell pellets were then resuspended in 1ml complete monolayer media (See table) and plated directly onto cell culture-treated plates.

### Establishment of primary human breast monolayer cultures

FACS-purified basal (CD45^−^/CD31^−^/CD49f^hi^/EpCAM^lo-med^) and luminal progenitor (CD45^−^/CD31^−^/CD49f^hi^/EpCAM^hi^) populations were pelleted by centrifugation at 350 g for 5 min. Cells were then resuspended in complete monolayer media (See table below) and seeded directly onto cell culture-treated plates or dishes, with a 30,000–50,000 cells/cm^2^ density. Media was refreshed in 24 hrs following confirmation of cell attachment and then every 1-3 days thereafter. Basal and luminal progenitor (LP) monolayers should attain distinct morphology and be passaged after 5-7 days.

### Passaging, freezing and thawing of primary human breast monolayer cultures

Monolayer cultures at 70-85% confluency were dissociated using pre-warmed Trypsin-EDTA (0.25%; STEMCELL Technologies, 07901) for ∼5 min. Cell suspensions were transferred to 15 mL conical tubes with room temperature DMEM/F12 (Wisent 319-085-CL) and centrifuged at 350 xg for 5 min, 4°C. The supernatant was aspirated, and the cell pellets were resuspended in 1 mL of complete monolayer media. Cells were seeded into new plates or dishes at 10,000–30,000 cells/cm^2^. Media was refreshed in 18-24 hrs following passaging and then every 1-2 days thereafter. In preparation for liquid nitrogen storage, cell pellets are resuspended in an in-house freezing media comprised of 70% complete monolayer media, 20% filtered fetal bovine serum (Gibco, 12483-020, Lot 2330093RP) and 10% DMSO (v/v), then aliquoted into 1.5 mL cryotubes. Cryopreserved monolayers were hand-thawed or warmed up in a 37°C water bath. Thawed cell suspensions were directly pipetted into pre-labelled plates or dishes with complete monolayer media at room temperature. Media was refreshed in 24 hrs following confirmation of cell attachment

### Media Formulation

- Base media: DMEM/F12 with L-Glutamine and 15 mM HEPES (319-085-CL, Wisent)
- Sterilize complete media using the 0.22 μm Steriflip filter (SCGP00525, Millipore Sigma)
- Use media within 2 weeks. Old media can be used to make freezing media for monolayers

| Component | Stock Concentration | Working Concentration | Volume for 50 mL of complete media |
| --- | --- | --- | --- |
| DMEM/F12 | - | - | 44.5 mL |
| FBS | - | 10% | 5 mL |
| Anti-Anti | 100x | 1x | 500 µL |
| Adenine | 0.1 M | 18 mM | 90 µL |
| Insulin | 4 mg/mL | 5 µg/mL | 63 µL |
| Hydrocortisone | 0.5 mg/mL | 0.5 µg/mL | 50 µL |
| Y-2763 | 10 mM | 10 µM | 50 µL |
| Cholera Toxin | 100 µg/ml | 2 ng/mL | 1 µL |
| hEGF | 100 µg/mL | 2 ng/mL | 1 µL |

### Establishment of on-top human mammary organoid cultures

Viably cryopreserved epithelium-enriched fractions (pellet A) from dissociated breast samples were thawed and digested with pre-warmed Trypsin-EDTA (0.25%; STEMCELL Technologies, 07901) for 3 min with trituration using a P1000 pipette. The cell suspension was quenched with 10 mL HF and passed through a 100-um filter to produce a suspension of cells and small cell fragments. The cell suspension was subsequently centrifuged at 300 g for 5 min, 4°C and the resulting cell pellet was resuspended in modified type 2 expansion media (T2 CGM; Dekkers et al. 2021) for cell counting by hemacytometer. Cells/cell fragments were seeded at an initial 20,000– 30,000 cells/cm^2^ in modified T2 CGM supplemented with 2% (v/v) type 2 reduced growth factor basement membrane extract (RGF BME; R&D Systems, 3533-010-02) onto cell culture-treated plates pre-coated with a thin layer of undiluted RGF BME as previously described (Lee et al. 2007). Media was refreshed every 2-3 days, where Y-2763 was removed from the modified T2 CGM 3 days after initial seeding. Modified T2 CGM is not supplemented with ovarian hormones, RSPO and Wnt ligands.

### Passaging, freezing and thawing of on-top patient-derived mammary organoid (PDMO) cultures

PDMOs were propagated every 7-12 days as small fragments using TrypLE Express (Thermofisher, 12605036) following published passaging practices (Rosenbluth et al. 2020, Dekkers et al. 2021). Established PDMOs were reseeded on top BME BME-coated plates at a 12,000– 18,000 cells/cm^2^ density. Dissociated cells/cell fragments from PDMOs were resuspended in Recovery Cell Culture Freezing Media (Thermofisher, 12648010) for liquid nitrogen storage. Cryopreserved PDMO samples were thawed in a 37°C water bath, washed with ice-cold Advanced DMEM/F12 (Thermofisher, 12634028) in 15 mL conical tubes and then pelleted by centrifugation at 300 g for 5 min, 4°C. The resulting cell pellets were resuspended in T2 CGM with Y-2763 for counting by hemacytometer or immediate seeding.

### Monolayer proliferation assay

Basal and LP monolayers were seeded in 6-well plates (8x10^4^ cells per well). After 24 hrs, the monolayers were rinsed gently with pre-warmed PBS, and complete monolayer media containing 1:2000 SPY505-DNA probe (Cytoskeleton Inc., CY-SC101) was added. After 2 hr incubation, the 6-well plates were placed into the BioSpa8 automated incubator (Agilent), where fluorescence images of 9 fixed ROIs from each well were captured every 2 hours by the Cytation 5 multimodal reader (Agilent) over 120 hrs. Image processing and cell nuclei counting were performed using the Gen5 software (3.14).

### Cell cycle analysis by imaging flow cytometry (Cytek AMINS ImageStream MkII)

Monolayer cells were incubated with 10mM Click-iT EdU for 2 hrs before being dissociated into a cell suspension with Trypsin-EDTA (0.25%; STEMCELL Technologies, 07901) as described prior. Cell suspensions in 5 mL round-bottom tubes were fixed with 4% PFA followed by permeabilization with 0.1% saponin & 1% BSA for 10 minutes at room temperature, respectively. 500 mL of freshly prepared 1X Click-iT reaction cocktail was mixed in with each sample tube and incubated for 30 min at room temperature, protected from light. 3 mL of 1X Click-iT® saponin-based permeabilization and wash reagent was used to wash each sample. Then, cells were stained with an antibody cocktail of the cell-surface mammary lineage markers, rat anti-human APC-CD49f (1:100; R&D Systems (clone GoH3), FAB13501A) and mouse anti-human PE-EpCAM (1:50; Biolegend (clone 9C4), 324206) prepared in PBS, for 30 min on ice. Following cell surface marker staining, cells were stained with 50 ng/mL (1:100,000) DAPI for 10 min at room temperature. After a final wash step, cell pellets were resuspended in 50-100 mL PBS and transferred to 1.5 mL Eppendorf tubes. Click-iT EdU kit reagents (Themofisher, C10425) were prepared according to manufacturer instructions. All centrifugation steps were performed at 350 xg for 5 min, at 4°C, before PFA fixation and 400g for 5 min, at 4°C after fixation. If not specified, all washing steps were performed with PBS.

### SA-beta-gal assay by imaging flow cytometry

Cellular senescence was assayed using CellEvent™ Senescence Green Flow Cytometry Assay Kit (Thermofisher, C10840). Cells were cultured in 6-well plates using the monolayer media and were irradiated using Best Theratronics Gammacell 40 exactor for the calculated time that can cause 10gy of g-irradiation. At the specified time post-irradiation, the cells were collected in 1.5mL tubes and washed once with 1X PBS. As described earlier, the cells were stained for cell surface lineage markers. Cells were washed with 1X PBS and spun down at 350 xg for 5 mins. Cells were fixed with 4% PFA by incubating with the PFA at 4°C for 15 minutes. Cells are washed again with 1X PBS and spun down to remove the buffer. The cell pellet is resuspended in 1:1000 diluted (in respective buffer) CellEvent™ Senescence Green Probe and incubated at 37 °C, without CO2, for 2 hours. After staining, the cells were washed with 1X PBS followed by nuclear staining using 1:10,000 diluted DAPI in 1X PBS. Cells were washed using 1X PBS before analysing by imaging flow cytometry.

### Protein lysate collection and western blotting

Cells cultured in 6-well treated plates were collected at specified time points post-irradiation. The media was removed, and cells were washed with 1X PBS to remove leftover media. 100mL of 2X SB was directly added on top of the cells and scraped to take the cell lysates. The lysates were collected into a 1.5ml tube and stored at -20° C. To run the samples on a gel, they were taken out and directly placed on a heat block at 96° C for 5 minutes for the first run and 1 minute for the further runs. They are then spun down at 13200 rpm for 1 minute. 12ml lysates were loaded to 4-12% gradient pre-cast gels kept in 1X MES SDS running buffer (Volume for the consecutive run is adjusted to match equal loading control levels based on densitometry). SDS gel was run at a constant 120V till the dye front reached the gel exit. The gel is transferred onto BIO-RAD Immun-Blot PVDF Membrane (0.2mm) using Invitrogen Mini-Blot module. Transfer buffer composition is mentioned in the table below

The transfer was done at 4°C for 2 hours using a constant current of 250mA. Further, the blot was blocked with 5% skimmed milk made in 1X PBST (1000ml 1X PBS+1ml Tween 20) for 1 hour at RT. Antibody dilutions were prepared using 5% BSA prepared in 1X PBS. Antibodies and concentrations used are listed in the table below. Blots were incubated overnight at 4°C with the antibody, and the next day, they were washed with 1X PBST 3 times. Secondary antibodies were prepared in 5% skim milk, and blots were incubated at RT for 2 hours. After 3 washes with 1X PBST, the blots incubated with fluorophore-conjugated secondary antibodies were imaged using LiCOR Odyssey CLx. If incubated with HRP-conjugated secondary antibodies, they were developed using ThermoFisher Supersignal West Femto Maximum Sensitivity Substrate (34095) and Bio-Rad ChemiDoc. Images are analyzed using ImageJ, and densitometric analysis was carried out and normalized to the untreated control samples in each experiment. The data are presented as fold change differences compared to their respective samples.

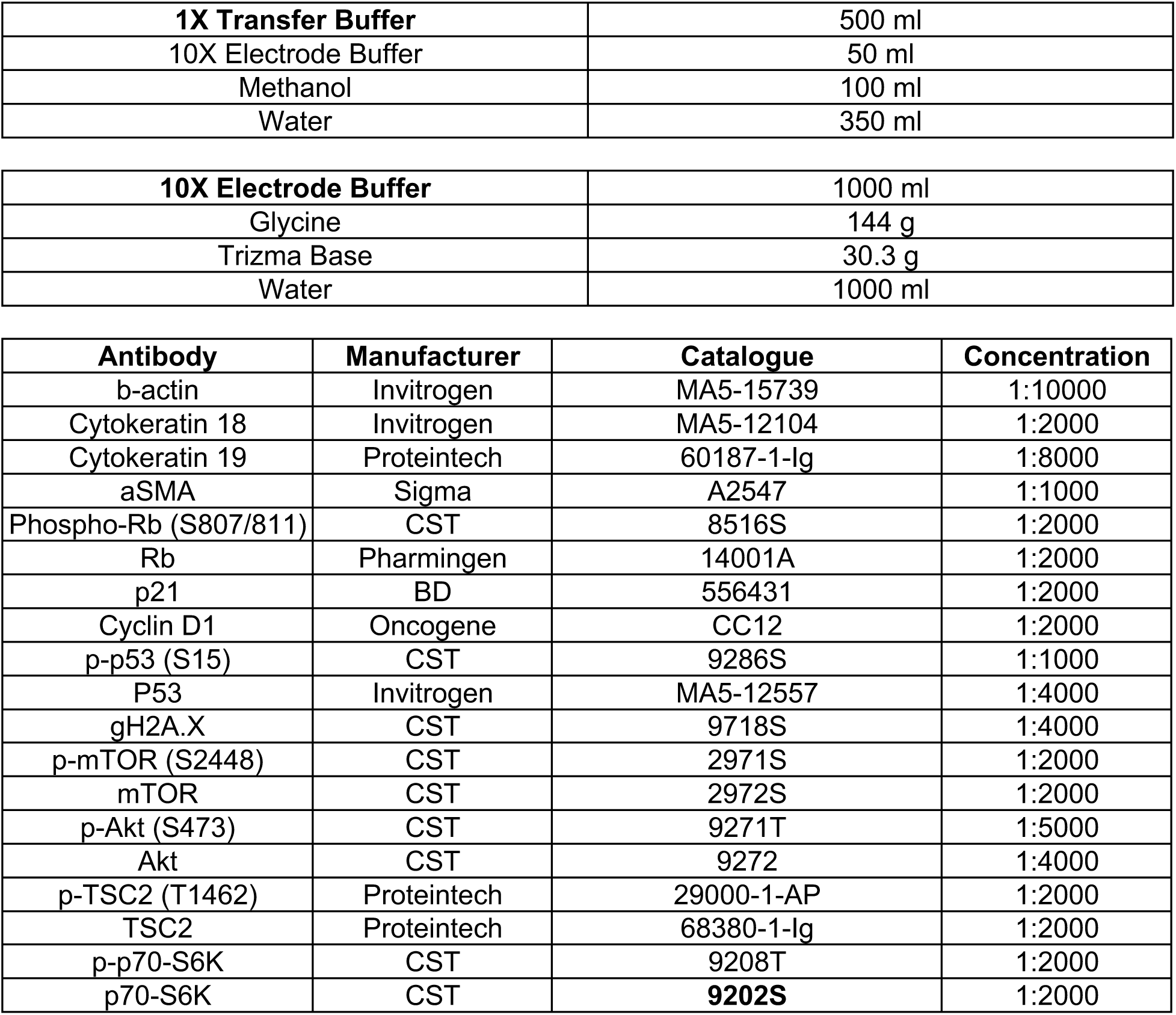

|  |  |
| --- | --- |
| **1X Transfer Buffer** | 500 ml |
| 10X Electrode Buffer | 50 ml |
| Methanol | 100 ml |
| Water | 350 ml |
| **10X Electrode Buffer** | 1000 ml |
| Glycine | 144 g |
| Trizma Base | 30.3 g |
| Water | 1000 ml |

| **Antibody** | **Manufacturer** | **Catalogue** | **Concentration** |
| --- | --- | --- | --- |
| b-actin | Invitrogen | MA5-15739 | 1:10000 |
| Cytokeratin 18 | Invitrogen | MA5-12104 | 1:2000 |
| Cytokeratin 19 | Proteintech | 60187-1-Ig | 1:8000 |
| aSMA | Sigma | A2547 | 1:1000 |
| Phospho-Rb (S807/811) | CST | 8516S | 1:2000 |
| Rb | Pharmingen | 14001A | 1:2000 |
| p21 | BD | 556431 | 1:2000 |
| Cyclin D1 | Oncogene | CC12 | 1:2000 |
| p-p53 (S15) | CST | 9286S | 1:1000 |
| P53 | Invitrogen | MA5-12557 | 1:4000 |
| gH2A.X | CST | 9718S | 1:4000 |
| p-mTOR (S2448) | CST | 2971S | 1:2000 |
| mTOR | CST | 2972S | 1:2000 |
| p-Akt (S473) | CST | 9271T | 1:5000 |
| Akt | CST | 9272 | 1:4000 |
| p-TSC2 (T1462) | Proteintech | 29000-1-AP | 1:2000 |
| TSC2 | Proteintech | 68380-1-Ig | 1:2000 |
| p-p70-S6K | CST | 9208T | 1:2000 |
| p70-S6K | CST | **9202S** | 1:2000 |

### Monolayer RNA extraction for bulk RNA-seq

Primary tissue-derived monolayer cells were cultured in 6-well cell culture-treated plate at 150x10^3^ cells per well density. 16 hours post seeding, the cells were irradiated along with the media using Best Theratronics Gammacell 40 Exactor a specific time that is validated to cause 10Gy of g-irradiation. 6 hours or 24 hours post irradiation, the media is removed from the plate, and cells are washed with 1X PBS and 250ml TRIzol was added. The cells were scraped and collected, and are all stored at -80° C. When all the samples were collected, the TRIzol lysates were thawed on ice and proceeded with the standard TRIzol-based RNA extraction protocol. After the RNA pellet is dried, they are dissolved in 15mL of Nuclease Free water. The RNA was confirmed to contain at least 10ng/ml RNA using Qubit RNA HS assay. They were shipped to Novogene for RNA sequencing. Libraries were prepared by Novogene Inc using ABClonal mRNA-seq library prep or NEBNext Ultra II RNA kits. Libraries were sequenced to a targeted 20 million paired end reads with Illumina NovaSeq X Plus.

### RNA-seq data processing

Adapter sequences were trimmed using Trimgalore v0.6.6 (CutAdapt v 4.8, with the following parameters: -q 25 --stringency 5 --length 20 --paired –a AGATCGGAAGAGCGTCGTGTAGGGAAAGAGTGTAGATCTCGGTGGTCGCCGTATCATT -a2 GATCGGAAGAGCACACGTCTGAACTCCAGTCACGGATGACTATCTCGTATGCCGTCTTCTGCTTG. Trimmed sequences were aligned with Spliced Transcripts Alignment to a Reference (STAR, v2.7.9a) with human reference genome release 46 (GRCh38.p14) with the default parameters for paired end reads. Transcripts were quantified using RNA-Seq by Expectation Maximization (RSEM, v1.3.0) with default parameters. Differential expression analyses at and between individual time points and in the time-series were performed using DESeq2 v1.44.0. Sample information was introduced as a covariate in the design formulae to correct for sample specific effects. Expression patterns of differentially expressed genes in the time series were clustered using the DEGreport R package (v1.40.1).

### Bulk RNA-seq data availability

All bulk RNA-seq raw files and processed result files acquired in this study will be publicly available from the Gene Expression Omnibus (https://www.ncbi.nlm.nih.gov/geo/).

### Proteomic sample preparation

Primary tissue-derived monolayer cells were cultured in 6-well cell culture-treated plate at 1.5x10^4^ cells per well density. 16 hours post seeding, the cells were irradiated along with the media using Best Theratronics Gammacell 40 Exactor a specific time that is validated to cause 10Gy of g-irradiation. At 6- and 24-hour time points post irradiation, the media is removed from the plate, and cells are washed with 1X PBS, and scrapped to collect as pellet.

The remaining buffer was removed by drying using SpeedVac Vacuum Concentrator (ThermoFisher). Pellets were resuspended in 50 μL of PBS with 2% (w/v) SDS and then were heated at 95 °C for 5 min using a heat block. Samples were sonicated (VialTweeter; Hielscher Ultrasonics, Teltow, Germany) by three ten-second pulses, set on ice for one minute, and then sonicated by three ten-second pulses. Samples were brought to 5 mM tris (TCEP) and allowed to reduce at 37 °C for 30 min at 1200 rpm on Thermomixer (Eppendorf). Samples were then brought to 10 mM iodoacetamide and incubated for alkylation at 37 °C for 30 min at 1200 rpm on Thermomixer. Samples were digested for 16 hr using SP3^53^ using 10 μL of prewashed magnetic particles (100 µg/µL of 1:1 SeraMag Hydrophilic:SeraMag Hydrophobic) and 1μg of Trypsin/Lys-C (Promega, V5072) for each sample. Peptides were de-salted using C18-based solid phase extraction.

### Mass spectrometry data acquisition

LC-MS/MS analysis was performed on an Orbitrap Eclipse MS (ThermoFisher) coupled to Neo Vanquish (ThermoFisher) chilled to 4 °C. Peptides were washed on pre-column (Acclaim™ PepMap™ 100 C18, ThermoFisher) with 60 μL of mobile phase A (0.1% FA in HPLC grade water) at 10 μL/min separated using a 250 nL.min flow rate over a 110 cm μPAC™ Neo HPLC Columns (COL-NANO110NEOB, ThermoFisher) ramping mobile phase B (0.1% FA, 80% HPLC grade acetonitrile in HPLC grade water) from 0% to 5% in 2 min, 5-35% in 134 min, 35% to 70% in 20 min interfaced online using an EASY-Spray™ source (ThermoFisher). The Orbitrap Eclipse MS was operated in data dependent acquisition mode using two 1.5 s cycles at different FAIMS settings (CVs of -45 and -60) with a full MS resolution of 240,000 with a full scan range of 375-1050 *m/z* with RF Lens at 60%, full MS AGC at 250%, and maximum inject time at 50 ms. MS/MS scans were recorded in the ion trap with 0.6 Th isolation window, 20 ms maximum injection time, with a scan range of 200-1400 *m/z* using Rapid scan rate. Ions for MS/MS were selected using monoisotopic peak detection, intensity threshold of 1,000, positive charge states of 2-5, 20 s dynamic exclusion, and then fragmented using HCD with 27.5% NCE.

### Mass spectrometry raw data analysis

Raw files were analyzed using FragPipe (v.20.0) using MSFragger^54,55^ (v.3.8) to search against a human proteome (Uniprot, 43,392 sequences, accessed 2023-02-08) – canonical plus isoforms. Default settings for LFQ workflow^56,57^ were applied using IonQuant^58^ (v.1.9.8) and Philosopher ^35^ (v.5.0.0) with the following modifications: Precursor and fragment mass tolerance were specified at -50 to 50 ppm and 0.15 Da, respectively; parameter optimization was disabled; MaxLFQ min ions was set to 1; MBR RT tolerance was set to 1 min, and MBR top runs was set to 10.

#### Mass spectrometry statistical analysis

All analysis was performed using R programming language (v.4.5.3) with Tidyverse pacakge (tidyverse_2.0.0) unless otherwise specified. The “MaxLFQ Intensity” columns were extracted from the “combined_protein.tsv” output file from FragPipe. Protein imputation was performed in two stages. First, proteins were split by lineage (e.g. basal and luminal) and then were filtered for presence in > 75% of samples. Each lineage was subsequently imputed with random forest algorithm using the MissForest package (missForest_1.6.1). Next, the original dataset was filtered for > 50% detection in basal or luminal samples. Missing values were first replaced by those calculated from the lineage-specific random forest imputation. Then, the remaining missing values were imputed using lower-tail imputation with a spread of 0.4 σ and a downshift of 2 σ.

#### Mass spectrometry data availability

All mass spectrometry raw files and processed result files acquired in this study will be publicly available from UCSD’s MassIVE database (ftp://massive.ucsd.edu).

#### Gene Set Enrichment Analyses

Rank scores for each gene from the differential expression analyses were calculated with the following formula: (sign of log2(fold change)) * (-1) * log10(p-value). The scores were used in pre-ranked Gene Set Enrichment Analyses (GSEA) with default parameters.

References

Trimgalore (cite this for adaptor trimming along with CutAdapt): https://github.com/FelixKrueger/TrimGalore

CutAdapt: https://journal.embnet.org/index.php/embnetjournal/article/view/200 STAR: PMID 23104886 https://pubmed.ncbi.nlm.nih.gov/23104886/

RSEM: PMID 21816040 https://pubmed.ncbi.nlm.nih.gov/21816040/ DESeq2: PMID 25516281 https://pubmed.ncbi.nlm.nih.gov/25516281/ Limma: PMID 25605792 https://pubmed.ncbi.nlm.nih.gov/25605792/

DEGreport: Pantano L (2024). DEGreport: Report of DEG analysis. R package version 1.40.1, <http://lpantano.github.io/DEGreport/>.

Human reference genome, from Gencode: PMID 30357393 https://pubmed.ncbi.nlm.nih.gov/30357393/

GSEA: PMID 16199517 https://pubmed.ncbi.nlm.nih.gov/16199517/ PMID 12808457 https://pubmed.ncbi.nlm.nih.gov/12808457/

#### Heatmap generation

Trimmed regularised log normalised counts data or log normalised proteomic intensity data were used for heatmap generation in pretty Heatmap (/pheatmap) in R studio using R version 4.3.0. Genelist of lineage heatmaps were compiled based on Mahendralingam *et al.* 2021.

#### Pathway Mapping

RNAseq counts data processed through the DEseq2 pipeline to compare each treatment timepoint with its respective control samples. The results were filtered for genes with log2FC>/= (+/-)0.5 and padj </=0.05. These genes were projected to KEGG pathways using *pathview* and *clusterprofiler* packages^60,61^. The generated maps were edited manually to improve visibility and clarity.

## Acknowledgements

This work is supported by funding from Canadian Institutes of Health Research (CIHR), Breast Cancer Canada, Canadian Cancer Society Research Institute, and the Terry Fox Research Institute PPG. R.K, S.M.H, and T.K. were supported through the Canadian Research Chair program. BZ is supported by a CIHR Graduate Research Scholarship (Doctoral) and O.D. is supported by an Ontario Graduate Scholarship. S.S. is supported by a TRIANGLE doctoral award. A.K is supported by a Princess Margaret Postdoctoral Fellowship and MW is supported by a CIHR Postdoctoral Fellowship. C.W.M was supported by a Banting Postdoctoral Fellowship. Graphical abstract and workflow schematics were created with BioRender.com. We would like to thank H.K.B and C.W.M for procurement and curation of the breast tissue cohort. Fluorescence activated cell sorting was performed using instruments from the Princess Margaret Flow Cytometry Facility at the Princess Margaret Cancer Centre.

## Author Contributions

A.K., B.Z., M.W., P.T. designed, performed experiments, and S.S., and O.D. contributed to data acquisition. M.W. acquired the mass spectrometry data using equipment and instruments in the T.K. lab. H.K.B., C.W.M. and H.F. provided support on breast tissue acquisition, processing and repository curation. A.K., B.Z., M.W. and K.H.Y. analyzed the data. A.K., B.Z and K.H.Y generated figures. A.K. and B.Z. wrote the manuscript, which all other authors edited and approved. R.K. and S.M.K. supervised the study.

---

**Figure S1.**
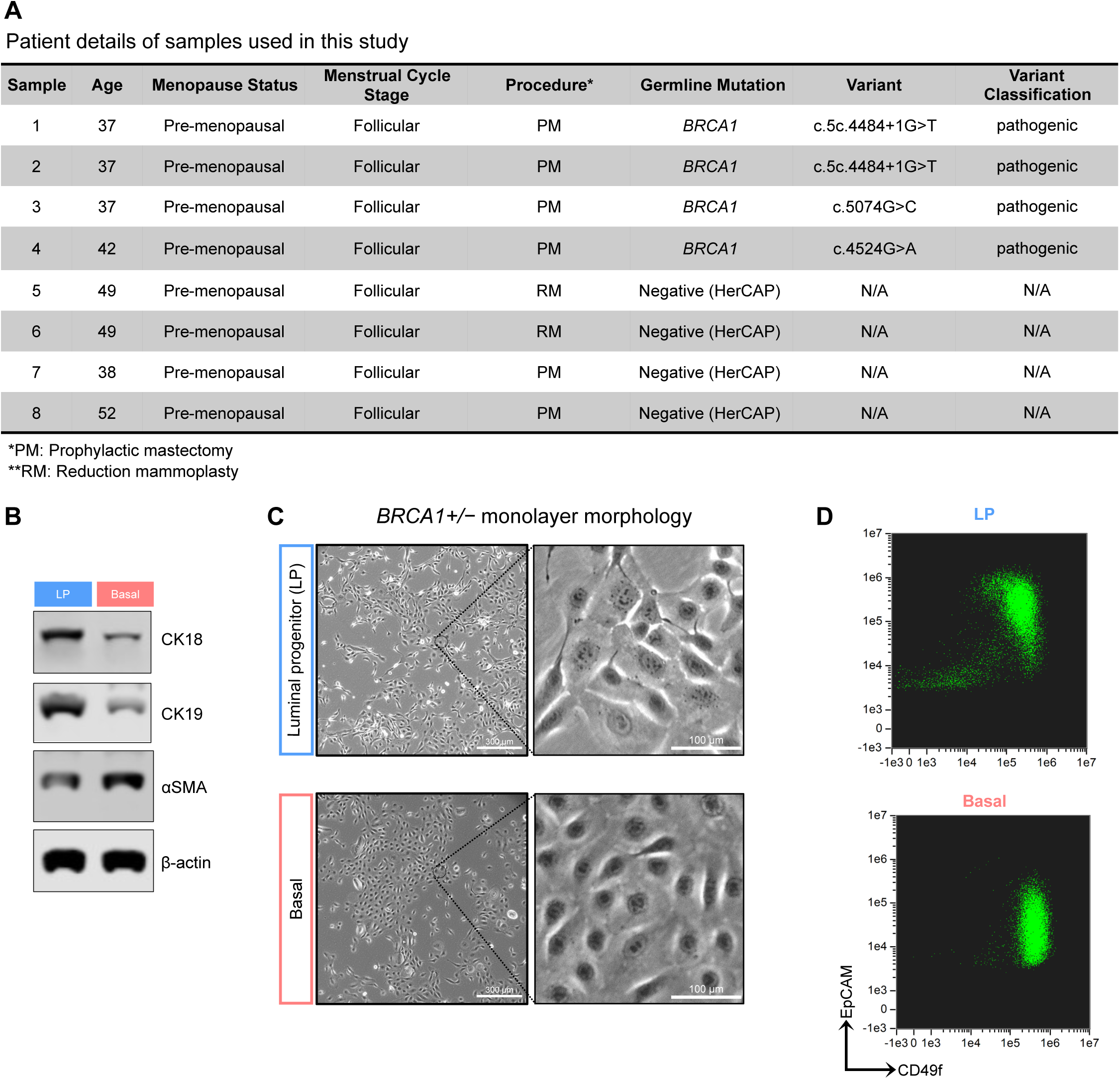
Lineage feature verification of monolayers from *BRCA1* mutation carrier and breast-derived organoid cultures. A) Details of the patient samples used in the study. B) Representative western blot images showing expression of lineage markers in monolayer cultures after 3 passages. C) Representative images of LP and basal monolayer cultures derived from *BRCA1* mutation carriers (Scale-300 mm) and their corresponding D) flow cytometry profile after 3 passages.

**Figure S2.**
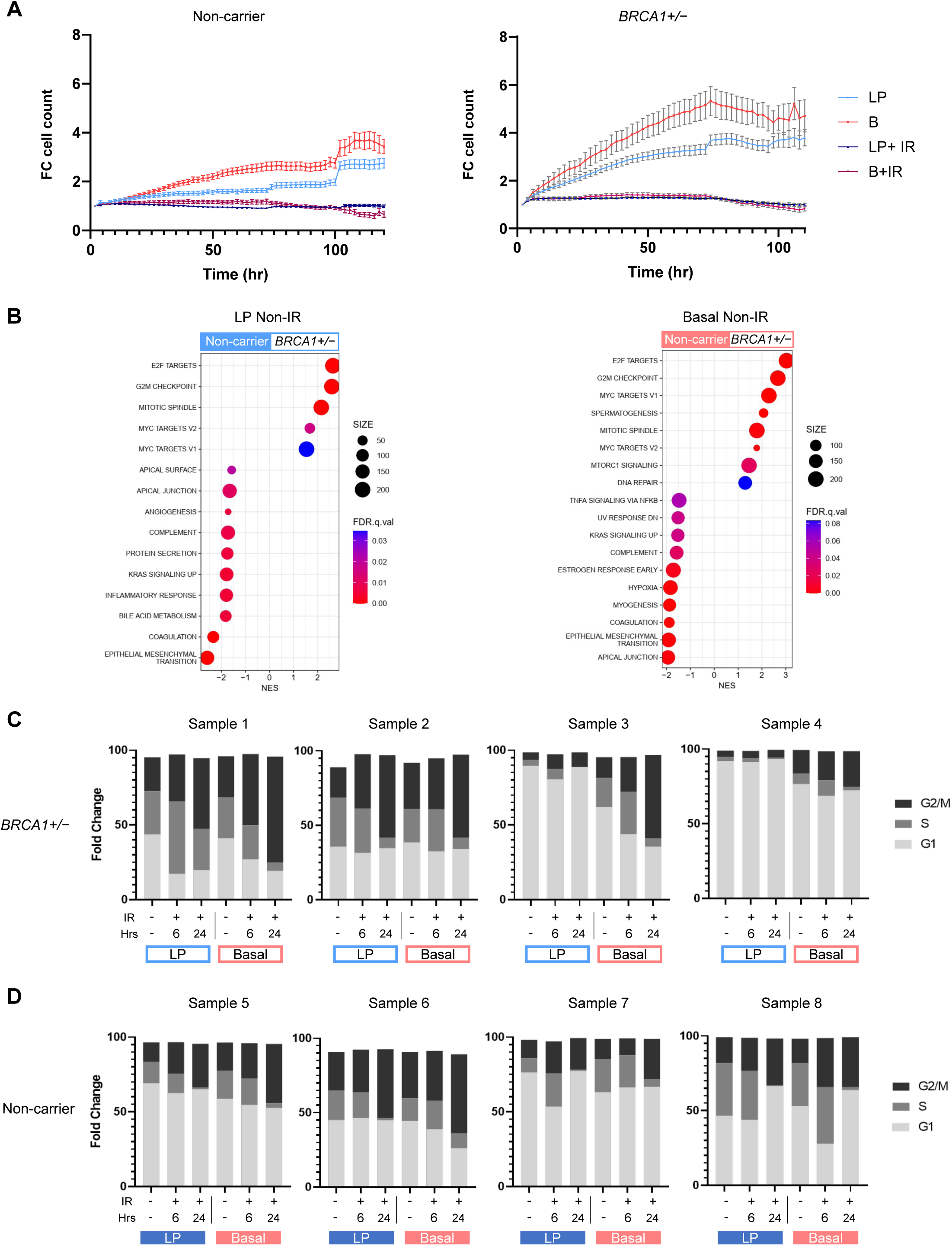
*BRCA1*+/− LP and basal cells have enhanced proliferative capacity. Proliferation (Fold change of nuclei count) of A) non-carrier and *BRCA1*+/− monolayers treated with or without irradiation (10 Gy) imaged every 2 hours over 120 hours. B) Transcriptomics-driven GSEA showing top 10 Hallmark pathways enriched in unirradiated *BRCA1*+/− LP and basal cells relative to corresponding non-carrier cells. Stacked bar plots showing cell cycle phase distributions in each biological replicate post irradiation of the C) *BRCA1*+/− and D) non carrier monolayer cultures.

**Figure S3.**
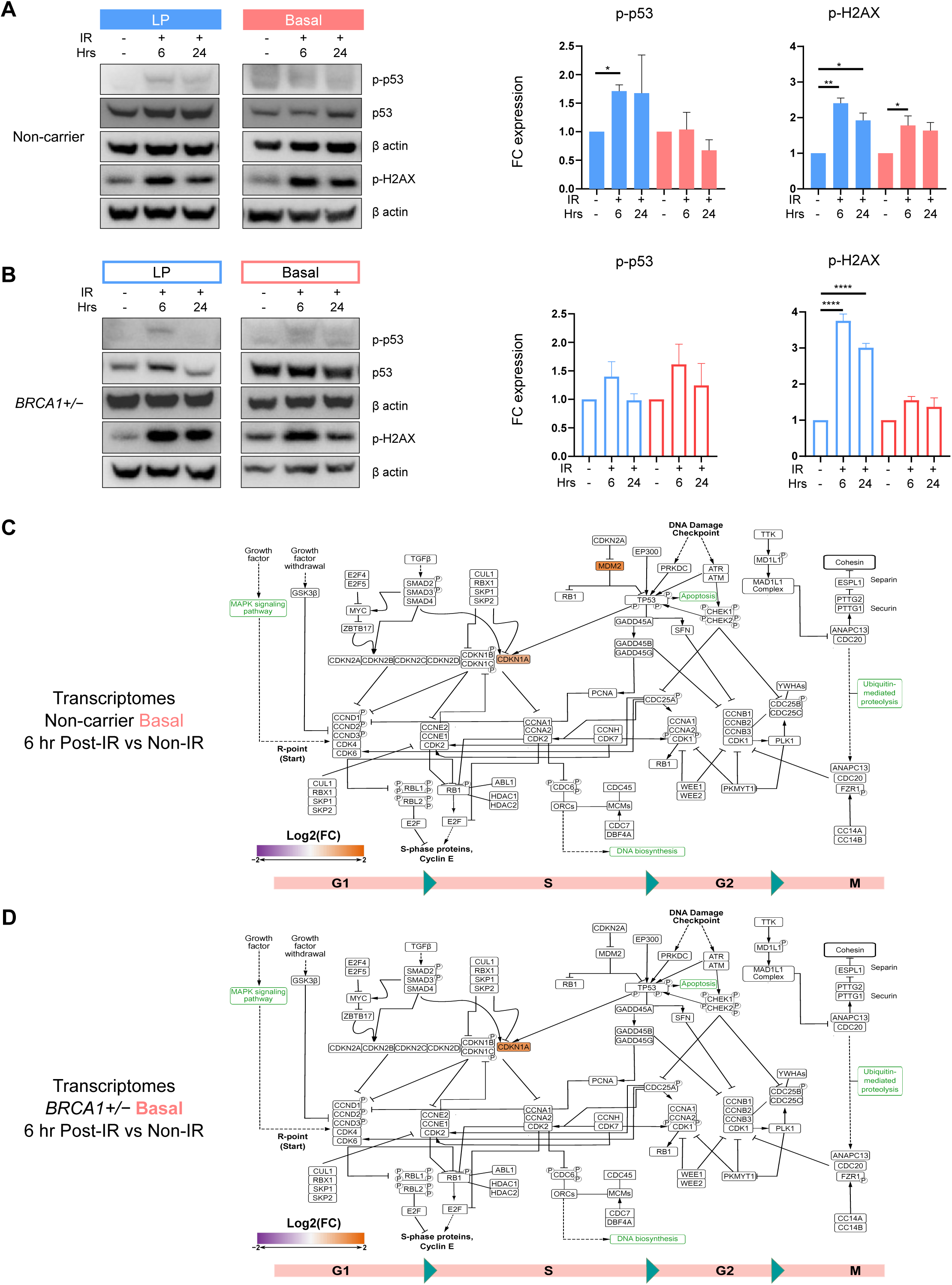
Irradiation-induced cell cycle regulation in basal cells is similar between *BRCA1* mutation carriers and non-carriers. Representative western blots probed for phosphor-p53, phospho-H2AX and β-actin along with corresponding densitometry quantification plotted as box plots of fold change in normalized expression relative to non-irradiated control in in A) non-carrier and B) *BRCA1*+/− monolayers (N=4, 1-way ANOVA and Holm-Sidak’s multiple comparison). KEGG cell cycle pathway diagram showing significantly altered transcripts with a FC of at least 1.5 relative to non-IR control for C) non-carrier and D) *BRCA1*+/− basal monolayers 6 hours post irradiation.

**Figure S4.**
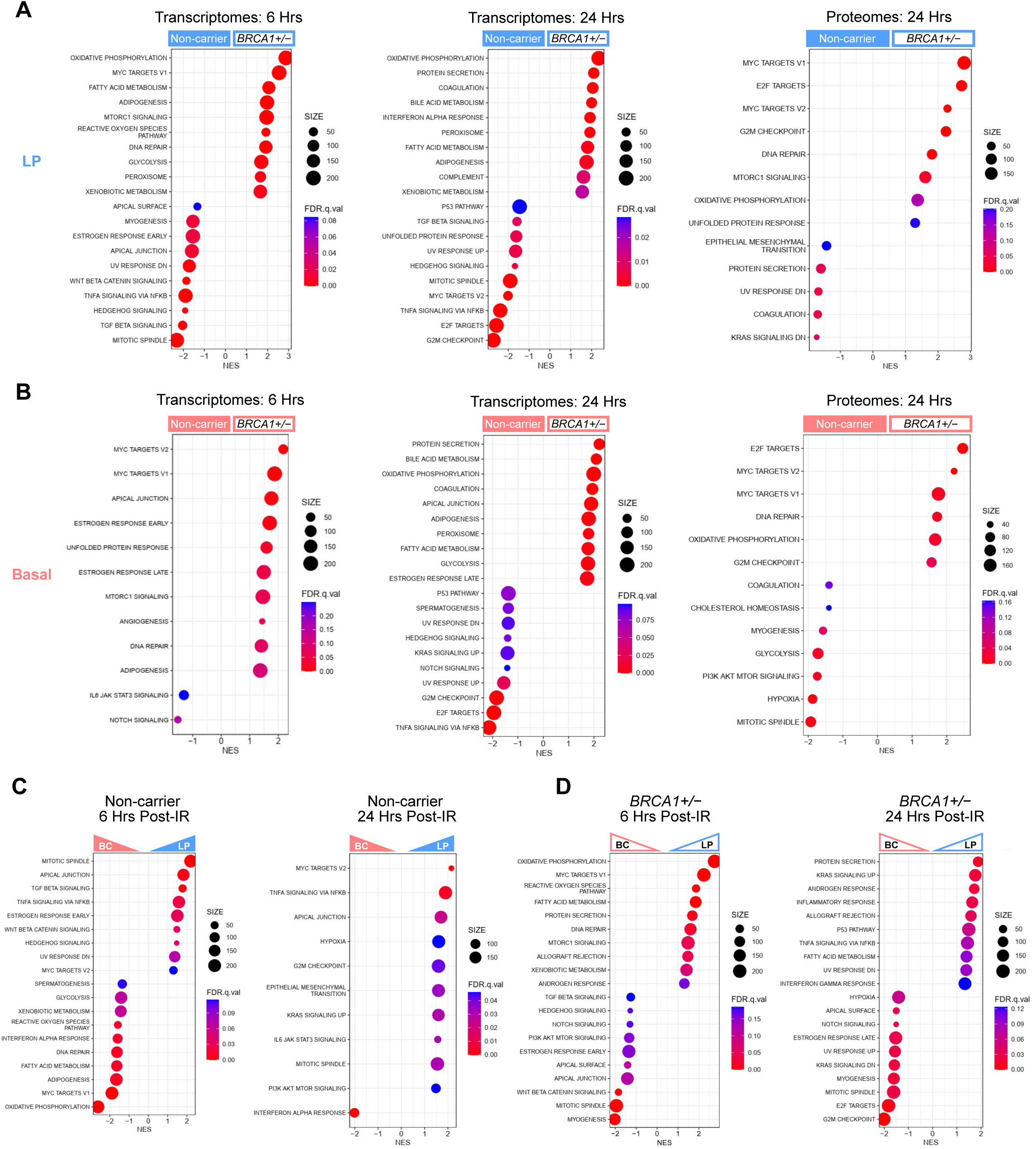
Omics-driven comparisons suggest deregulated metabolic and senescence-associated signalling in BRCA+/- cells. GSEA showing top ten differentially enriched Hallmark gene sets from transcriptomics at 6- and 24-hours post IR and from proteomics at 24-hours post IR, for non-carriers compared to *BRCA1*+/− in A) LP and B) basal monolayers. GSEA showing top ten differentially enriched Hallmark gene sets from transcriptomics between lineages (LP vs Basal) at 6- and 24-hours post IR in C) non-carriers and D) *BRCA1*+/− monolayers.

